# Harm to others acts as a cingulate dependent negative reinforcer in rat

**DOI:** 10.1101/808949

**Authors:** Julen Hernandez-Lallement, Augustine Triumph Attah, Valeria Gazzola, Christian Keysers

## Abstract

Empathy, the ability to share another individual’s emotional state and/or experience, has been suggested to be a source of prosocial motivation by making actions that harm others aversive. The neural underpinnings and evolution of such harm aversion remain poorly understood. Here, we characterize an animal model of harm aversion in which a rat can choose between two levers providing equal amounts of food, but one additionally delivering a footshock to a neighboring rat. We find that independently of sex and familiarity, rats reduce their usage of the preferred lever when it causes harm to a conspecific, displaying an individually varying degree of harm aversion. Prior experience with pain increases this effect. In additional experiments, we show that rats reduce the usage of the harm-inducing lever when it delivers twice, but not thrice the number of pellets than the non-preferred lever, setting boundaries on the magnitude of harm aversion. Finally, we show that pharmacological deactivation of the anterior cingulate cortex, a region we have shown to be essential for emotional contagion, reduces harm aversion, while leaving behavioral flexibility unaffected. This model of harm aversion might help shed light onto the neural basis of psychiatric disorders characterized by reduced harm aversion, including psychopathy and conduct disorders with reduced empathy, and provide an assay for the development of pharmacological treatments of such disorders.

## Introduction

Learning to avoid actions that harm others is an important aspect of human development [1], and callousness to other’s harm is a hallmark of antisocial psychiatric disorders including psychopathy and conduct disorder with reduced empathy [2]. What could motivate humans and other animals to refrain from harming others? An influential theory posits that vicarious emotions (i.e. emotions felt by a witness, in the stead of the witnessed individual) including emotional contagion and empathy, trigger harm aversion [3]. Put simply: harming other people is unpleasant, because we vicariously share the pain we inflict. Accordingly, it has been argued that psychiatric disorders characterized by anti-social behavior [2,4] might stem from malfunctioning or biased vicarious emotions [5,6].

An increasing number of studies show that mice and rats display affective reactions to the pain of others observed as increased freezing and changes in pain sensitivity while witnessing the pain of others [7–11] and when re-exposed to cues associated with the other’s pain [12,13]. Recent studies in rats identified emotional mirror neurons involved in the processing of pain also activated while witnessing other’s distress [14,15] in the anterior cingulate cortex (ACC; area 24 in particular). Reducing activity in the ACC reduces emotional contagion [9,15]. However, in these paradigms, the observing rat is not the cause of the witnessed pain. Whether vicarious activity in area 24 is associated with harm aversion thus remains unclear.

Inspired by classic studies [16,17] here we refine a paradigm to study instrumental harm aversion in rats. A rat called the “*actor*” can press one of two levers for sucrose pellets. After a baseline phase revealing the rat’s preference for one of the levers, we pair this preferred lever with a shock to a second rat (“*victim*”), located in an adjacent compartment (Figure 1). We then measure how much actors switch away from the preferred lever as a behavioral index of harm aversion.

**Figure 1.**
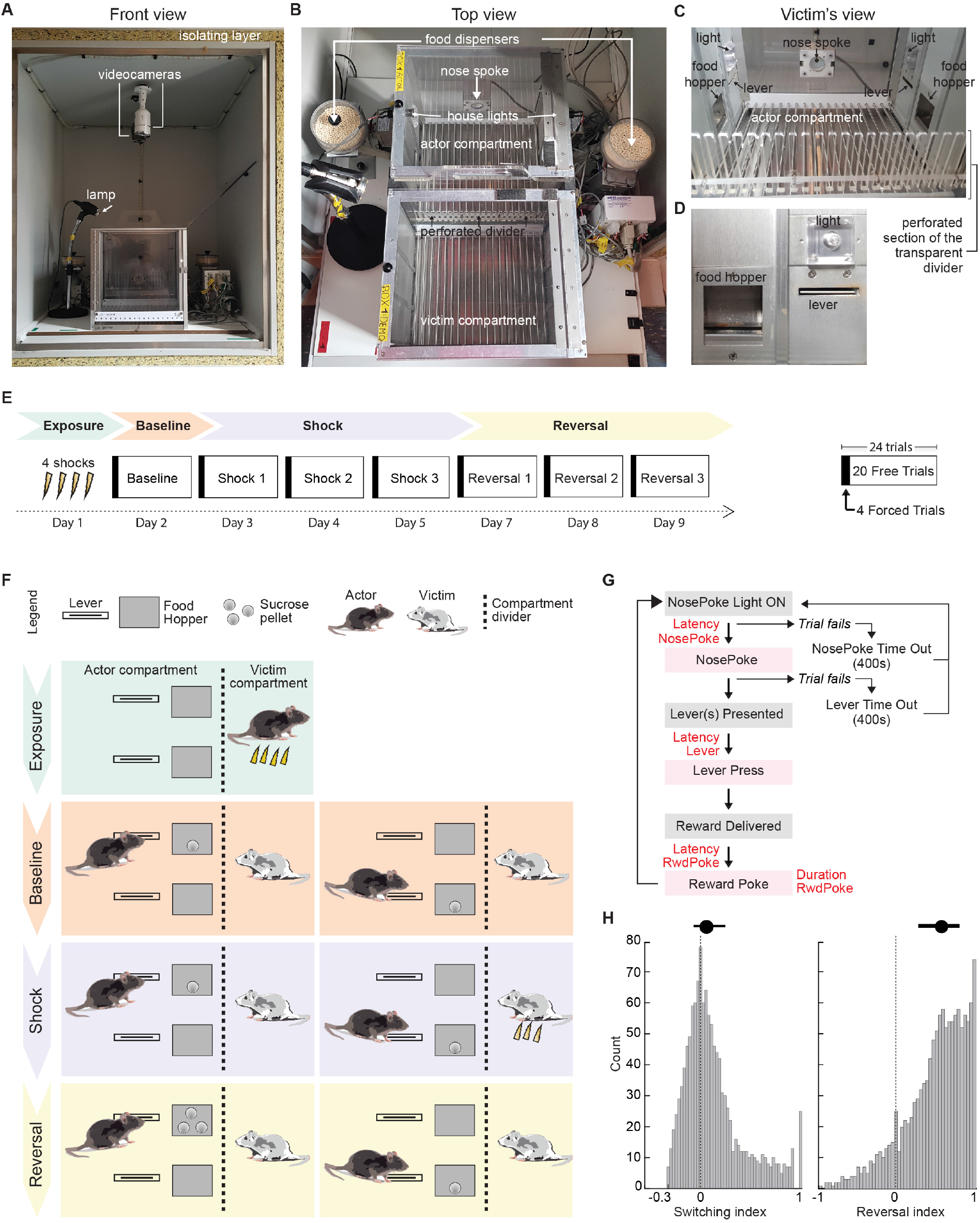
Experimental Procedures. **(A)** Photo of the cabinet in which the experimental set-up was isolated. **(B)** Top view of the two-compartment set-up. **(C)** View of the actor’s compartment through the perforated transparent divider as seen from the victim’s compartment. **(D)** Close-up of the left wall with one of the two levers and food hoppers available to the actor. **(E)** Timeline of the procedures. **(F)** Design of exposure, baseline, shock and reversal sessions. Note that the reversal session is only present in some conditions (Table 1, 2). During exposure, the actor receives 4 shocks alone in the victim compartment. During Baseline, pressing one lever (left column) or the other (right column) leads to one pellet for the actor. The lever preferred during baseline then additionally triggers a shock to the victim during the 3 days of the Shock phase. During reversal, shocks are no longer delivered, but the non-preferred lever now leads to 3 pellets. **(G)** Trial structure for training, baseline, shock and reversal sessions. Grey coloring denotes events and purple, the rat’s responses. **(H)** Simulated distribution for switching (left) and reversal indexes (right) given their respective equations and uniform distributions of preferences. Dots and lines above the distributions are the median and the 25 and 75 % percentile.

We show (i) both male and female rats switch significantly away from the shock-delivering lever; (ii) this is stronger in shock-pre-exposed actors; and (iii) deactivating the ACC reduces this effect. By altering the timing of shock delivery, we show contingency between lever-pressing and shock-delivery is essential. By varying the reward value of the levers, we show rats switch from an easier to a harder lever and from one that provides two pellets to one that provides one pellet to prevent harm to another. However, rats were unwilling to switch from a lever that provides three pellets to one that provides one pellet. We additionally report and explore substantial individual differences in switching across Sprague-Dawley rats.

**Table 1.** Core experimental conditions. For each of the four experimental conditions (columns), the table specifies who got electrical shocks during the exposure, baseline and shock sessions. A ‘/’ indicates that the victim was not present during the exposure session. Sample size (N) reflects the number of actors included in the behavioral analyses. **♂**:male. **♀**:female.

|  | ContingentHarm<br>N = 24 (12♂, 12♀) |  | NoHarm<br>N = 14, all ♂ |  | RandomHarm<br>N = 8, all ♂ |  |
| --- | --- | --- | --- | --- | --- | --- |
|  | Shock to Actor | Shock to Victim | Shock to Actor | Shock to Victim | Shock to Actor | Shock to Victim |
| Exposure | Yes | / | Yes | / | Yes | / |
| Baseline | No | No | No | No | No | No |
| Shock | No | Yes | No | No | No | Yes |
| Reversal | No | No | No | No | No | No |

**Table 2.**
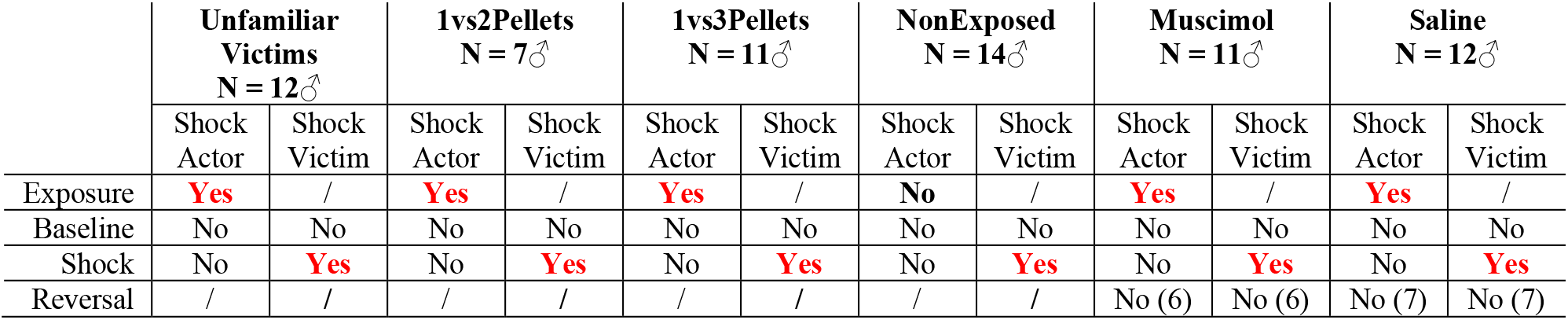
Paradigm contingencies for additional conditions. Convention as in Table 1. ‘/’ either indicates that the victim was not present or that the session was not run. Values in brackets for the Muscimol and Saline Reversal condition indicate the subsample for which this condition was run.

|  | Unfamiliar Victims<br>N = 12♂ |  | 1vs2Pellets<br>N = 7♂ |  | 1vs3Pellets<br>N = 11♂ |  | NonExposed<br>N = 14♂ |  | Muscimol<br>N = 11♂ |  | Saline<br>N = 12♂ |  |
| --- | --- | --- | --- | --- | --- | --- | --- | --- | --- | --- | --- | --- |
|  | Shock Actor | Shock Victim | Shock Actor | Shock Victim | Shock Actor | Shock Victim | Shock Actor | Shock Victim | Shock Actor | Shock Victim | Shock Actor | Shock Victim |
| Exposure | Yes | / | Yes | / | Yes | / | No | / | Yes | / | Yes | / |
| Baseline | No | No | No | No | No | No | No | No | No | No | No | No |
| Shock | No | Yes | No | Yes | No | Yes | No | Yes | No | Yes | No | Yes |
| Reversal | / | / | / | / | / | / | / | / | No (6) | No (6) | No (7) | No (7) |

## Results

### Rats switch away from a lever that triggers foot-shocks to a conspecific

We first compare the behavior of rats in three main conditions: ContingentHarm, NoHarm and RandomHarm (Table 1; Figure 1E,F). In all three conditions, an actor was trained to press one of two levers for one sucrose pellet in the actor compartment (Figure 1A-D). One lever was twice as hard to press as the other, with the harder-to-press side randomized across animals. After initial training alone, all actors were exposed to 4 foot-shocks (Exposure; Figure 1E,F) in the adjacent victim’s compartment to maximize emotional contagion [7,14,18]. Actors were then placed back into their actor compartment and performed 24 trials of lever pressing with their cage-mate in the victim compartment (Baseline).

These 24 trials started with 4 forced trials (2 for each lever; pseudo-randomized) to force actors to sample both options, followed by 20 free choice trials to measure baseline lever-preference (Figure 1E). In the ContingentHarm condition, on the 3 days following Baseline (Shock1, Shock2 and Shock3 sessions; Figure 1E), the actor performed 24 trials of the same task each day (4 forced + 20 free choice), similar to baseline trials except that pressing the lever preferred during Baseline triggered a foot-shock (0.8mA, 1s) to the victim in the adjacent compartment. In this condition, we had two groups: male pairs (ContingentHarm **♂**) and female pairs (ContingentHarm **♀**). We compared this condition against a NoHarm control condition, in which pressing either lever never delivered a shock to the victim. Finally, to explore the importance of action-outcome contingency, we include a RandomHarm condition. For the RandomHarm condition, we identified the 8 actors from the ContingentHarm condition (from all 24 animals) that showed the strongest switching away from the shock-lever. For each, we recorded the sequence of shock and no-shock trials to the victim. In the RandomHarm condition, each victim then received the sequence of shocks from one of the switchers from the ContingentHarm condition, independently of what lever was pressed by the actor, and with the shocks delayed randomly by 3-8s after actors exited the food receptacle to break the action-outcome contingency.

We then compared the change in preference from baseline to shock sessions across conditions. A 4 group (ContingentHarm♂, ContingentHarm♀, NoHarm♂ and RandomHarm♂) * 4 session (Baseline, Shock1, Shock2, Shock3) repeated measures ANOVA revealed a significant effect of session (*F*_(3,123)_ = 7.34, *p* < .001, *η*^2^ = .15) and a significant session*group interaction (*F*_(9,123)_ = 1.93, *p* = .05, *η*^2^ = .12). To follow up on these findings, we first concentrated on male actors, for which we have three groups (ContingentHarm**♂**; NoHarm♂; RandomHarms♂). At Baseline, the groups showed similar preference for the future shock lever (Figure 2A), and a Bayesian ANOVA revealed a BF_10_ = .32, showing evidence for the absence of group effect during baseline. ContingentHarm male actors then showed a significant decrease in shock-lever pressing from baseline to all shock sessions (dark green in Figure 2A), with their preference for the shock-lever lower than the NoHarm and RandomHarm control groups in all three shock sessions (pairwise comparisons; Figure 2A). Actors in the male ContingentHarm group thus shifted significantly away from a lever that causes shocks to a conspecific. Importantly, the change is not simply due to the distress of the victim (which was matched across ContingentHarm and the RandomHarm group), but to the contingency between the actions of the actor and the reactions of the victim.

**Figure 2.**
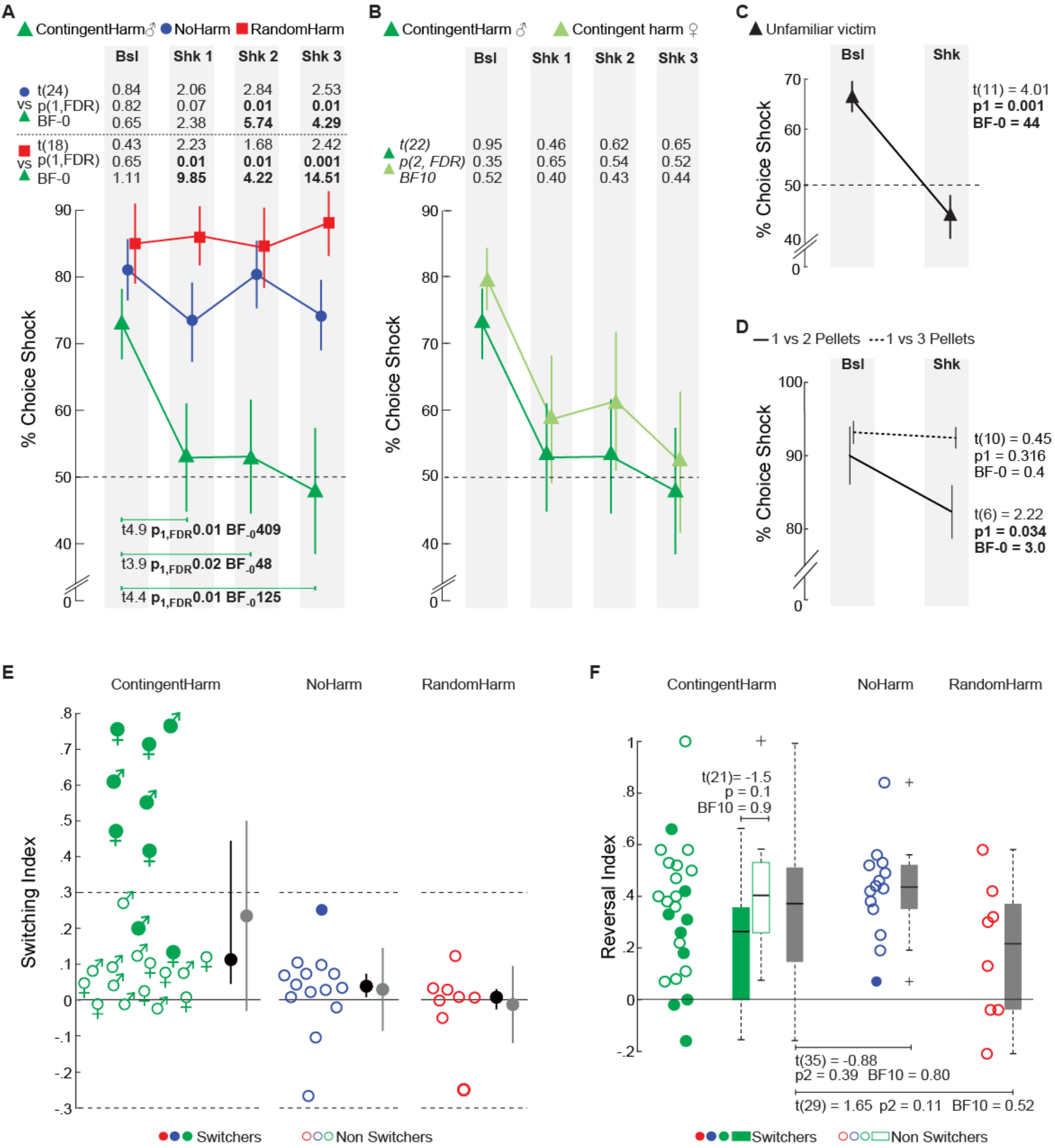
Harm aversion in rats. **(A)** Percent choice for shock lever across sessions (Baseline, Shock1, Shock2 and Shock3) and conditions (ContingentHarm, NoHarm and RandomHarm). The numbers above the graph specifie the t-statistic (t), one-tailed FDR corrected p (p1,FDR) and the one-tail Bayes factor (BF-0) for the comparisons between the two conditions indicated leftmost, separately for each session. In the lowest part of the graph, statistics for pair-wise comparison between each shock session and the baseline. Values in bold are significant. Bsl=Baseline. Shk1-3=Shock sessions 1-3. Gray rectangles on background help visually discriminate the sessions. Data are presented as mean +/- s.e.m. **(B)** Same as in (A) but comparing female and male ContigentHarm groups, and reporting two-tailed t-test (p2,FDR) and two-tailed Bayes Factor (BF10). **(C)** Average percent shock lever presses during baseline and shock sessions (i.e. average over 2 baseline sessions and 2 shock sessions) for the actors paired with unfamiliar victims. Conventions as in A and B. **(D)** Percent shock-lever presses for baseline and average of 3 shock sessions for 1 versus 2 pellets and 1 versus 3 pellets. **(E)** Switching index for the different conditions. Black dot and line: distribution’s median and the 25 and 75 % percentile values. Grey dot and line: distribution’s mean and standard deviation. **(F)** Reversal index for the different conditions. Boxplot of each distribution show the median (black bar) and outliers (crosses). For the ContingentHarm condition the Boxplots have been computed separately for the switchers (green filling), the non-switchers (green contour) and the whole group (gray filling). For the NoHarm and RandomHarm, only the group results are presented (gray filling) because there are insufficient switchers. See also Figure S1 for related results, and data.xlsx for the choice data.

We found no evidence of sex effects on harm aversion, when quantified as the change in preference across session (Figure 2B; *session*gender: F*_(3,63)_ = .21, *p* = .89, *η*^2^ = .01). The Bayes factor for including a gender*session was *BF*_10_ = .20, providing evidence for the absence of a sex difference on the change of lever preference. For all subsequent analyses looking at the change of preference across session, we thus pool males and females into one single ContingentHarm condition (N = 24 actors). The Bayes factor for including a main effect of gender, however, was anecdotal (*BF*_10_=0.44).

### Familiarity with the victim is not necessary for switching

During the piloting phase of the paradigm, we tested actors that were unfamiliar with their victims taken from unrelated cages (Unfamiliar Victims; Table 2). Actors showed a significant decrease from baseline preference levels also for shocks to these Unfamiliar Victims (paired one-tail t-test; Figure 2C). This effect was comparable to the one observed in the ContingentHarm animals of our main experiment (*session*condition; F*_(1,34)_ = .20, *p* = .66, *η*^2^ = 0.006). The Bayes factor for including a familiarity*session was *BF*_10_ = .80, which is inconclusive. Accordingly, familiarity is not necessary for reducing lever preference, in line with data showing that rats freeze even when an unfamiliar conspecific gets a shock [19] and free an unfamiliar trapped rat [20]. However we cannot exclude that familiarity may have a subtle effect on the magnitude of switching, as shown in mice [21–25]. It is important to note that during the piloting phase of the experiment, actors in the Unfamiliar Victims condition were also exposed to shocks prior to the experiment, but in another context rather than in the victim compartment (see supplementary materials). Hence, this condition shows that the switch in preference observed in the ContingentHarm condition is not solely due to contextual fear formed during the exposure session in the victim’s compartment.

### Switching is modulated by cost

To test whether actors would give up food to avoid another’s distress we tested new groups of rats (1vs2Pellets and 1vs3Pellets; Table 2) where levers required the same effort, but differed in rewards from the beginning of the baseline session. In the 1vs2 condition, the shock lever provided n = 2 pellets, while the no-shock lever provided n = 1 pellet. Actors decreased their preference for the 1vs2 option upon association with a shock (*paired one-tailed t-test;* Figure 2D, solid line). To explore whether this switching differed from the ContingentHarm, we computed a repeated measures ANOVA using ContingentHarm and 1vs2Pellets conditions with 4 sessions each. We found a highly significant main effect of session *(F*_(3,84)_ = 7.03, *p* < .001, *η*^2^= .20), but no interaction (*F*_(3,84)_ = 1.82, p = .15). The Bayes factor for condition*session was *BF*_10_ = 2.96, showing evidence leaning in favor of a difference across the conditions. Hence, rats are willing to forgo one sucrose pellet to avoid the victim’s distress, but the effect tends to be slightly reduced compared to a difference in effort. The 1vs3 pellet condition, where the levers led to n = 1 vs n =3 pellets (Figure 2D, dotted line) did not show a significant decrease of preference from baseline (*paired one-tailed t-test*), and a rmANOVA (2 Groups_[ContigentHarm, 1vs2Pellets]_ x 4 Sessions_[Bsl, Shk1, Shk2, Shk3]_) shows the effect was significantly smaller than in the ContingentHarm condition (*F*_(3,96)_=5.46, *p*=0.002) suggesting that harm aversion may not be strong enough to counteract high costs. Because delays are known to steeply devalue rewards in rodents, in a pilot study using a reduced number of animals (N = 5♂), rats were tested in a task where the same 1pellet reward was delayed by 2s when delivered through the no-shock lever (as opposed to immediately delivered for the shock lever). In this condition we did not find significant switching (*paired one-tailed t-test baseline vs shock; t*_(4)_ = 1.5, *p* = .21), but the Bayesian analysis shows that with such a small group-size this finding remains inconclusive (*BF*_10_ = 0.8). This suggests that 2 s delays may be too costly to generate a large effect that can be seen in a small group.

### Rats show substantial individual differences in switching

To explore individual differences, we computed a switching index (SI; see methods) which reflects a change in preference from baseline, normalized to obtain a score with positive values reflecting a switch away from the shock lever, with SI=1 representing the maximum possible switch given an individual’s baseline preference. Some animals showed a robust preference change in shock sessions, others remained indifferent to the victim’s distress, but none showed strong negative scores (Figure 2E), which would indicate an attraction to the lever that causes a shock. A permutation test where actual choices were randomly reassigned to the Baseline *vs* Shock sessions (see methods) revealed that n = 9 actors (i.e. 38%) in the ContingentHarm condition (n=4 males and n=5 females) showed a significant switch (at *p* < .05; green colored in circles in Figure 2E; hereafter referred to as *‘Switchers’*). A binomial test showed that 9 out of 24 switchers is not explained by chance (*binomial, N= 24, alpha =*. 05, *p* = 10^−6^). These *Switcher* animals found across males and females, showed a decrease between [25%-80%] from Baseline. Switching rates were within chance level in the NoHarm (N = 1 significant *Switcher*, blue colored in circle; *binomial, p = .51*) and absent in the RandomHarm condition (N = 0 significant *Switchers*). A χ² square test revealed that the ContingentHarm condition had significantly more *Switchers* than the NoHarm condition (*χ² = 4.20, p = .04*) and the RandomHarm condition (*χ² = 4.17, p = .04*).

### Behavioral correlates of individual differences in switching

To further explore individual variability, we examined factors that may have been associated with the switching index.

#### Flexibility

at the end of the Shock sessions, we identified which lever was less preferred, and baited it with 3 pellets to see how strongly rats would change their preference for rewards (Reversal session; Figure 1E,F; Table 1,2). We computed individual Reversal indexes (RI; see methods; Figure 2F) which quantified the strength of reversal from the last shock session across the three successive Reversal sessions. ContingentHarm animals did not show significant differences in RI from the NoHarm (Figure 2F) and RandomHarm animals (Figure 2F). S*witchers* and *Non-Switcher* animals showed comparable RI in the ContingentHarm condition (Figure 2F), suggesting that non-switchers switch as much as switchers for rewards but not for shocks to others.

#### Attentional Markers

we examined the latencies to enter the food hopper after pressing the shock or no-shock lever, and the time spent in the food hopper (Figure 3A). For all statistical analyses on latencies, we transformed latencies using a log-transform, to ensure approximate normality as verified using the Shapiro-Wilk test. We found that upon delivering the shock to the victim in the first Shock session, *Switchers* (solid lines in Figure 3A) delayed their entry to the food hopper and shortened the time spent in it (*Session*_[*Baseline, Shock1, Shock2, Sochk3*]_\**Trial*_[*shock lever vs no-shock lever*]_\**Type*_[*switchers vs non-switchers*];_ *log latency F*_(3,154)_= 4.50, *p* = .005; *log duration F*_(3,154)_= 3.34, *p* = .02), providing behavioral evidence that the harm to the other captured attention and competed against food-seeking. This was not true in *Non-Switchers* (dotted lines in Figure 3A), and there was a significant correlation between changes in latencies and switching (Figure 3B; *r* = .78, *p* < .001). Lever differences in latency and duration were not observed in NoHarm and RandomHarm conditions (Figure S2).

**Figure 3.**
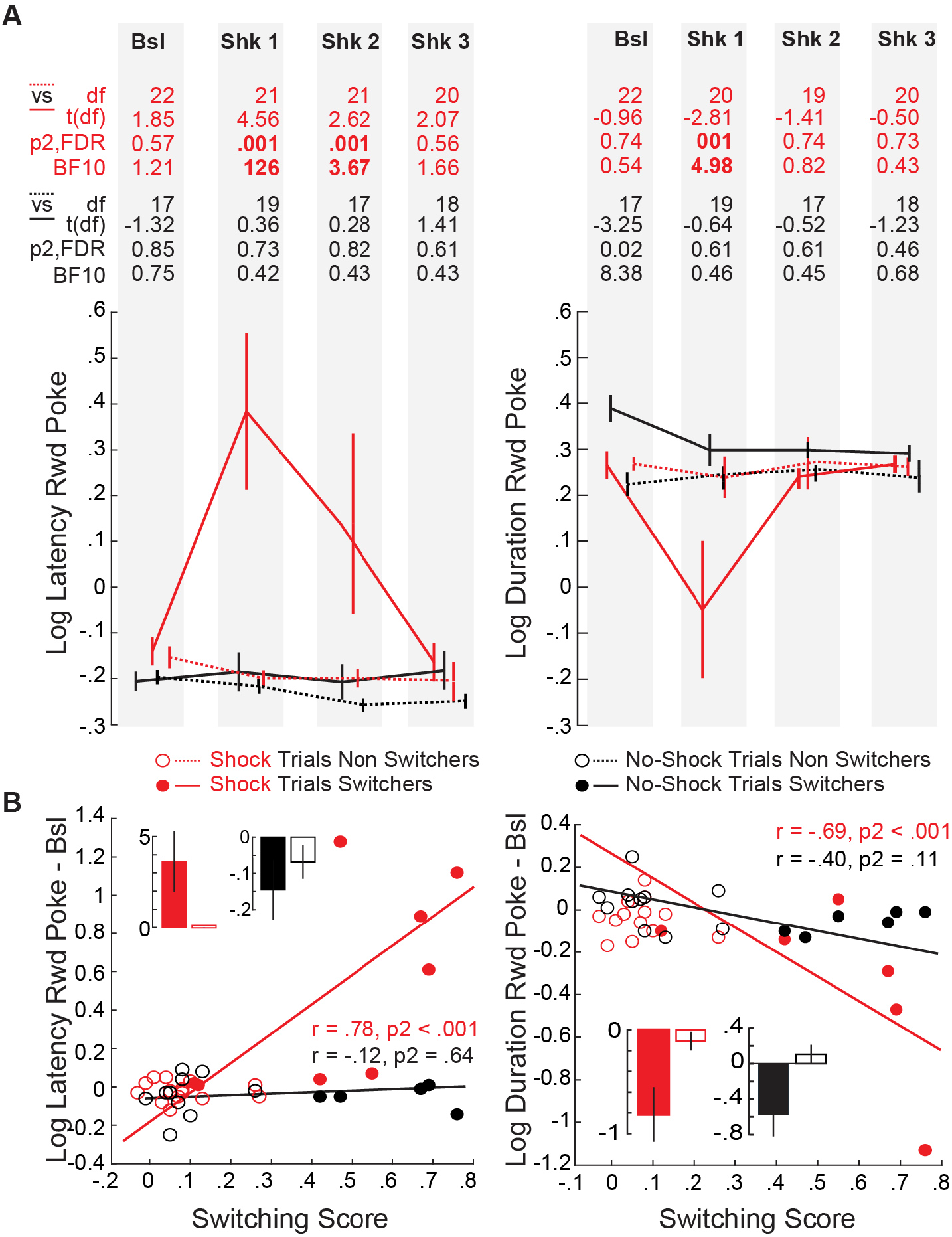
Behavioral correlates of switching. **(A)**. Log transformed reward poke latency (left) and log transformed poke duration (right), separately for switcher and non-switchers, and shock and no-shock trials in the ContingentHarm condition. **(B)** Correlation between switching scores and the change from baseline in log transformed reward poke latency (left) and log transformed duration (right). <u>Bar plots</u>: Mean raw (i.e. not log transformed) changes from baseline for reward poke latency (left) and duration (right) per trial type. Data is mean +/- s.e.m. p_1_ and p_2_ indicate one-and two-tailed testing, respectively. FDR: False Discovery Rate correction over 4 sessions. Bsl: Baseline. Shk: Shock sessions 1-3. df: degree of freedom, which are lower in sessions in which some animals never chose the no-shock or the shock lever. BF: Bayes Factor. See Figure S2 for similar data for the NoShock and RandomShock conditions.

### Exposure to foot-shocks potentiates switching

We explored whether switching is dependent on the intensity of the prior foot-shock experience [7,14,18]. During the Exposure sessions, animals increased freezing (one way ANOVA; *F*_(4,119)_= 24.36, *p* < .001) (Figure 4A), with animals showing more freezing during Exposure showing higher switching in the harm aversion paradigm (red in Figure 4B). A similar trend was observed in actors tested with Unfamiliar Victims (black in Figure 4B). To further test the importance of prior exposure, we tested a new group as in the ContingentHarm condition except that the actors received no shocks during the exposure session (NonExposed, Table 2). We observed no significant effect of session on lever-preference in NonExposed animals (Figure 4C, *one way ANOVA*, *F*_(4,119)_ = 0.56, *p* = .65, *BF*_10_=0.37), and a rmANOVA using both ContingentHarm and Non-Exposed conditions revealed a significant session (*F*_(3,55)_ = 8.71, *p* < .001, *η*^2^ = .20) and session*condition interaction (*F*_(3,105)_ = 3.51, *p* = .018, *η*^2^ = .09). While baseline preference levels were comparable across both conditions, ContingentHarm animals showed significantly lower preference for the shock lever during the first shock session compared to the NonExposed condition (Figure 4C; pairwise comparisons). This difference becomes less significant in the subsequent shock sessions. Interestingly, 2 out of 14 NonExposed animals showed significant decrease from baseline (Figure 4D). Together, this shows (i) that prior fear experience primes rats to a higher sensitivity to other’s pain but (ii) some rats do not need prior footshock experience to show harm aversion with other prior experiences perhaps generalizing to the observation of footshocks.

**Figure 4.**
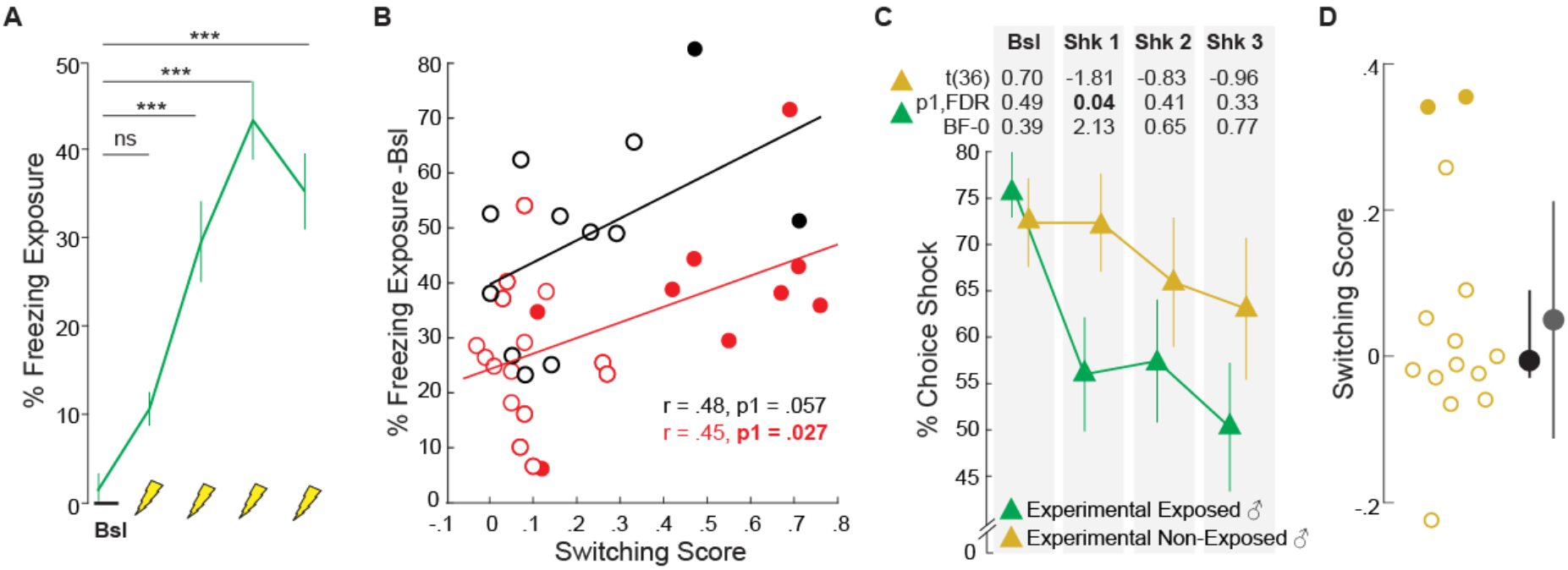
Shock exposure potentiates sensitivity to other’s distress. **(A)** Percentage freezing during the shock exposure session. Yellow lightening bolts indicate the four shocks, with percent freezing then quantifying the percentage freezing in the following intershock interval. Bsl: Baseline. **(B)** Correlation between the change in freezing levels relative to baseline during exposure and switching scores for the ContingentHarm group (red) and the Unfamiliar Victim group (black). Filled circles represent switchers, empty circles, the non-switchers. **(C)** Percent shock lever presses in ContingentHarm (**green**) and Non-Exposed conditions (**yellow**) across sessions. **(D)** Switching score distribution for Non-Exposed animals. Conventions as in (B). Black dot and line are the distribution’s median and the 25 and 75 % percentile values. Grey dot and line are the distribution’s mean and standard deviation. Data is mean +/- s.e.m. * *p* < .05, ** *p* < .01, ns= non significant, BF = Bayes Factor. p_1_ and p_2_ indicate one-tailed and two-tailed testing, respectively. FDR: p values corrected using False Discovery Rate for 4 sessions. See data.xlsx for the data that went into C.

### The ACC is necessary for harm aversion in rats

Several studies performed in humans [26–29] and rats [9,14,15,18] suggest the ACC (including area 24a and b, [30]) is recruited during the observation of distress, and maps other’s pain onto one’s own pain circuitry. To test whether the ACC is necessary for the switching in our paradigm, we infused muscimol bilaterally in the ACC in a group of rats (Muscimol; Table 2) prior to Baseline and Shock sessions and compared the choice allocation to a saline infused group (Saline; Figure 5A; Table 2). Infusions were centered at +1.8mm from the bregma, and had an anterio-posterior spread of [+1.95mm; +1.45mm] from bregma (Muscimol group: N = 9 out of 11; M = 1.76mm; SD = .26; Saline group: N = 9 out of 12, M = 1.84; SD = 0.26; Figure 5B), confirming that area 24a (approximately 0.6mm dorsal to corpus callosum) and area 24b were targeted. The infusion spared midcingulate areas 24’, located closer to the bregma [30,31], as well as deeper area 33 and the cingulum, located postero-ventral to most infusions [32]).

**Figure 5.**
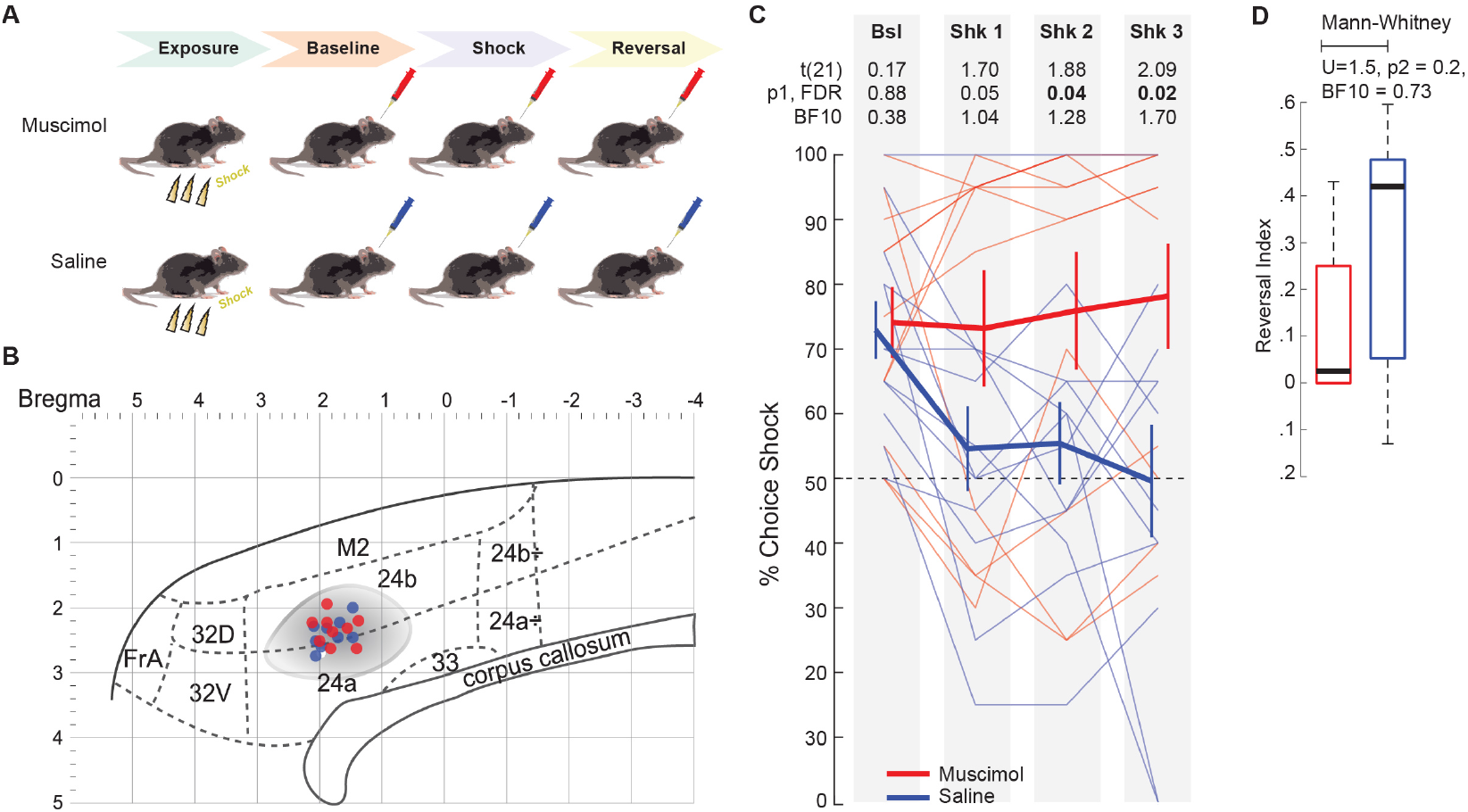
The ACC is necessary for switching. **(A)** Experimental procedures. **(B)** Estimated anterior-posterior and dorso-ventral coordinates of the infusions on a sagittal representation of the medial surface of the rat brain based on [30,33] for Muscimol (red) and Saline (blue) animals. Each dot is the average of the coordinates of the right and left canula tip location in histological slices. **Grey shading** estimate of likely spread based on a combination of published data [45] and estimates from our own lab based on similar injections of fluorescent muscimol. **(C)** Individual (lighter lines) and group (thicker lines) preferences for shock lever across sessions for Saline (**blue**) and Muscimol animals (**red**) **(D)** Reversal score for Muscimol and Saline groups did not differ. Data is mean +/- s.e.m. Significant numbers in bold. BF = Bayes Factor. p_1_ and p_2_ indicate one-tailed and two-tailed testing, respectively. pFDR=p value corrected by false discovery rate for 4 sessions. Also see data.xlsx for the choice data in C and D.

A 2 condition x 4 session rmANOVA revealed a trend for an effect of session (*F*_(3,60)_ = 2.32, *p* = .08, *η*^2^ = 0.10) and a significant session*condition interaction (*F*_(3,60)_ = 3.33, *p* = .025, *η*^2^ = 0.14). While baseline preferences were comparable between Muscimol and Saline, preference for the shock lever were significantly higher for the Muscimol than Saline infused animals in Shock sessions (Figure 5C)

To test whether muscimol reduced behavioral flexibility, we started including reversal sessions in later animals (Saline N=7 and Muscimol N=6; Table 2) by baiting the lever least preferred at the end of the shock session with 3 pellets, while removing shocks and continuing the injection of the respective drug. In this small sample, RIs were not significantly different between Muscimol and Saline conditions (Figure 5D), but the Bayesian analysis suggest a larger sample is necessary to exclude small effects on flexibility. If we repeat the ANOVA on the N=7+6 animals for which we performed reversal session and did not differ significantly on RI, we still find that switching is reduced after ACC deactivation (*2 condition*_[Muscimol, Saline]_ *x 4 session*_[Bsl,Shk1,Shk2,Shk3]_ *interaction*, *F*(3,33) = 5.6, *p* = .003).

## Discussion

By characterizing the willingness of rats to reduce the use of a lever that triggers pain to another conspecific, we provide evidence that rats display a contingency-dependent harm aversion, which we show to be mediated by ACC activity. Harm aversion is robust, and we replicate the effect in 5 separate groups of animals (male ContingentHarm, female ContingentHarm, Unfamiliar Victims, Saline and 1vs2 pellets) The tendency not to harm the conspecific was true whether the difference between the levers was twofold in effort or reward magnitude. However, the willingness to switch is no longer significant when the difference in value is too high (3 vs 1 pellet).

In the difference in reward magnitude experiments (1vs2 or 1vs3 pellets), switching entailed a reduction in reward value, because at baseline animals always preferred the 2 or 3 pellet levers over the 1 pellet lever. For the effort experiments, as observed in previous studies [17], approximately 25% of the animals actually preferred the hard to the easy lever in baseline sessions. For those, we then associated the hard lever with shocks, and switching involved a *reduction* in effort. We therefore interpret our data generally as suggesting that rats are willing to switch to a less-preferred lever to avoid shocks to others.

Our data provides a number of additional insights. First, we find sex does not modulate harm aversion. This is in apparent contrast to a small number of studies that reported sex effects on vicarious responses in mice [23,34] and rats [35,36], pointing towards a growing awareness that the specific output behavior measured can dramatically alter sex differences [37]. Second, the pain of the victim needs to be contingent on the actor’s actions: the same number of shocks to the victim without contingency (RandomHarm condition does not change lever-preference. Third, prior exposure to foot-shocks increases the behavioral effects of witnessed harm, in line with an increasing number of studies showing that vicarious freezing is increased by prior exposure [7,14,18].

The human literature has introduced a distinction between helping motivated by self-regarding personal distress vs. other-regarding sympathy [38]. That switching is linked to prior pain experience is compatible with the notion that, as for many humans [38], actors avoided the shock lever to reduce personal distress: the victim’s distress cues could reactivate aversive engrams from their own past pain-experience, which is aversive enough to act as a negative reinforcer. In this view, harm-aversion may not primarily be an altruistic motive to prevent pain to another rat, but a more selfish motive to avoid an unpleasant personal state triggered by the signals emitted by the other rat - a less noble but perhaps equally effective motive.

In accord with the original study of Greene [17], we found strong individual differences across our rats, with a subset showing moderate to strong switching, while the remaining animals did not. The exact proportion we find (38%) is probably dependent on many experimental details, but variability was a consistent finding in our data. While non-switchers were behaviorally flexible (as shown in the reversal session) and showed some degree of attention towards the victim (spending more time close to the victim on shock trials), their reaction time data shows the pain of the victim did not distract them from their pursuit of food as much. Such individual variability in our paradigm could be valuable to shed light on the origin of individual variance in human harm aversion, particularly given that rodent models of the kind of disrespect for other people’s wellbeing encountered in human antisocial behavior are so far lacking [39].

Finally, we found that deactivating the ACC reduces switching. This demonstrates the sensitivity of our paradigm to reveal the involvement of brain regions in harm aversion, Importantly, recent studies have suggested that both in rats and humans the pain felt by a conspecific is mapped onto our own pain representation through emotional mirror neurons, located within the ACC [14,15,40]. Our deactivation data finally provide evidence that this region is important to prevent harm to others.

## STAR Methods

### Methods

#### Subjects

Sprague Dawley rats were ordered from Janvier Labs (France). Rats were separated in different groups to test different parameters of the harm aversion paradigm (Table 1 & 2). Except for one group specifically testing females (Table 1), all animals were males. Rats were socially housed in groups of four same-sex individuals (SPF, type III cages with sawdust), in a temperature- (22-24 °C) and humidity-controlled (55% relative humidity) animal facility, on a reversed 12:12 light:dark cycle (lights off at 07:00). Water was provided ad libitum. Upon start of the training phase, a food deprivation schedule was implemented to maintain animals at 85% of their free-feeding body weight, which was monitored daily. Animals were pre-fed with 66% of their daily food intake 2h before the start of harm aversion testing (to prevent that high hunger masks harm aversion). All tests were done in the dark phase of the animals’ circadian rhythm, between 08:30 and 13:00. All animals were 30 days old at arrival, and started harm aversion testing at 55 days of age. Male and female rats weighed on average 302.4g (SD = 96.5) and 240.8g (SD = 14.1) at the start of the experiment. All experimental procedures were approved by the Centrale Commissie Dierproeven of the Netherlands (AVD801002015105) and by the welfare body of the Netherlands institute for Neuroscience (IVD, protocol number NIN171105-181103). Studies were conducted in strict accordance with the European Community’s Council Directive (86/609/EEC).

#### Sample size calculation

##### Behavioral experiment

one experiment performed in rats reported that changes in (n[rats] = 10) preferences contingent on caused conspecific distress was significant at *p* = .01[17]. Based on z-score tables (http://www.z-table.com/), we estimated that an equivalent z-score = 2.29, which outputted an effect size r = .72 (*r* = *zscore*/(*sqrt*(*N*)). For an effect size r = .72, an 80% power, α = 0.05, we computed a required sample size of N = 8 actors per condition. We thus aimed for groups of at least 8 actors in all conditions, and sometimes used larger samples to increase sensitivity. We supplement the frequentist approach with Bayesian analysis whenever no significant *p*-values were found to further inform the interpretation of negative findings.

##### Pharmacology experiment

given that the decrease in vicarious freezing in ACC deactivated animals corresponds to an effect size r = 1.22 [15], we used the same effect size for sample size calculations in the pharmacological experiment. For an effect size r = 1.22, an 80% power, α = 0.05, we computed a required sample size of N = 12 animals. Accounting for ~10% of dropout (e.g. surgery casualties, wrong target of the implanted cannula), we expect N = 14 actors for each muscimol and saline condition, summing to a total of N = 28 animals.

#### Experimental setup

The experimental setup (Figure 1A-D) consisted of two skinner boxes (ENV-008CT; Med Associates, Inc.) fused into one single setup in our Mechatronics department. The separation walls between the compartments (i.e., skinner boxes) were replaced by a single perforated Plexiglas wall that allowed the transmission of auditory, olfactory and visual information between compartments. The compartment’s floor consisted of a stainless grid floor (ENV-005). One of the compartments’ floor (the victim’s compartment) was linked to a stimulus scrambler that allow the delivery of a foot-shock (ENV-414S). In the actor’s compartment, one nose poke unit (ENV-114BM) was installed on the wall opposite to the divider (Figure 1B-C). Two retractable levers (ENV-112CM) equipped with a stimulus light above each of them (ENV-221M) and a food hopper (ENV-200R2MA) beside each of them, were placed on the lateral walls of the actor’s compartment (Figure 1C-D). Both levers were equidistant from the nose poke unit. Two food dispensers (ENV-203-45IR) placed outside on each side the actor’s compartment allowed the delivery of sucrose reward pellets for correct lever presses in the food hoppers (Figure 1B). Finally, two house lights (ENV-215M) were placed close to the top of the box above each dispenser (Figure 1A-B). House lights were turned ON before and after the session, which indicated that no operant items could be operated by the animals. The setups were placed in sound attenuating cabinets (Drefa, The Netherlands; Figure 1A). Rats performed all tasks in the dark. Infrared cameras (one per compartment; 600 TVL 6 mm; Sygonix) were used to record the rats’ behavior.

#### Experimental procedures

##### Acclimation, handling and habituation

On the day of arrival, rats were randomly housed in groups of four and were allowed to acclimate for four days to the colony room. The animals’ tails were marked from 1 to 4 to identify the animals in each cage. Numbering was random to avoid that animals less anxious would be labelled first in each cage. After 4 days rats were handled during the dark phase of the cycle for 5 minutes for five days. Individuals #1 & #3 of each cage (1 and 3 stripes) were assigned to the actor role and individuals # 2 & #4 (2 and 4 stripes) were assigned to the victim role. Within each cage, two pairs were formed always following the same pattern: ACT#1 and VIC#2 formed one pair and ACT#3 and VIC#4 the second. The two members of a pair were always tested together, thus ensuring familiarity. There was only one exception to this rule: the Unfamiliar Victim condition that specifically investigated the effect of unfamiliarity within a dyad. On the fifth day of handling, animals were transported within their home cage to the experimental room, where the rats were placed in their compartment for 5 minutes then weighted and placed back in their home cage. Between each rat, the compartment of the skinner box was cleaned with 70% ethanol. This was done consistently throughout each phase of the whole experiment. To facilitate the acquisition of reward-driven lever press, sucrose pellets (n=3) were placed in the reward on the first day of training. From this day on, the animals’ food intake was reduced to bring animals to 85% of their free feeding body weight, which was monitored on a daily basis.

##### Training

On the session following habituation to the setup, the actors were shaped to press levers to obtain food. To do so, animals went through 3 steps of training.

*Step 1:* Actors were placed in their compartment, with house light ON. The session was started by the experimenter using an adjacent computer, which was indicated to the animal by turning off the house light. Both levers were constantly presented and could be pressed by the animal. Either lever required 30cN to be operated. No times out were used, so as to maximize the change for animals to accidentally press one of the levers. Either lever press led to the delivery of n = 3 sucrose pellets in the adjacent reward hopper. Animals had a maximum session length of 20min, and could perform a maximum of 50 trials per session. Hence, animals could perform the session in less than 20min if they performed all 50 trials. A correct trial was a trial were the animal pressed one of the two levers. Promotion criteria to Step 2 was performing at least 35 lever presses out of the possible 50 (70%) within the 20min.

*Step 2:* Actors went through 5 blocks of 10 trials, leading to a total of 50 trials per session. Each block started with 4 forced trials (only one lever extended, 2 trials per lever, randomized order) followed by 6 free choice trials (both levers extended). Each trial started with the extension of the lever(s), and actors had 20s to press one of the two extended levers. Animals could press only one lever per trial, i.e., pressing one lever led to the retraction of the opposite lever and the delivery of reward in the adjacent receptacle. Either levers required 30cN to be operated. Lever press led to the delivery of n = 1 sucrose pellet as well as the activation of the light cue located above the lever for 1s. Pellets were delivered 1s after lever press. Animals had a maximum of 30min to perform all 50 trials. The 4 forced trials of each block were mandatory, and timeout (20s without pressing) lead to the repetition of that trial. For free choice trial, a timeout lead to the retraction of the levers and switching to the next trials. Promotion criteria was to press a lever within 20s on at least 70% of the 30 free choice trials, i.e. on 21 trials.

*Step 3:* Here, a nose poke was required in order to activate the levers and initiate the trial. In the first session, the nose-poke light was turned on for 50s, and a 10ms nose poke within those 50s was enough to trigger the extension of the levers. Animals then had 20s to press a lever, and perform a trial correctly. Performing at least 70% correct trials of the 30 free choice trials was used as a criterium to increase the required nose poke duration to 200ms, and the duration of the nose-poke illumination to 20s. Again, if 70% of the free choice trials are performed correctly, animals arrived at the final nose poke duration of 400ms to ensured that animals initiated the trial deliberately and not by accident. A timeout of 20s was implemented for nose poke and lever press, which ensured that animals performed trials in a rapid fashion. Hence, animals performed a minimum of 150 trials (i.e., three levels on nose poke duration) in this step. For the conditions testing magnitude (1vs2Pellets and 1vs3Pellets) and delay (0vs2Seconds), the levers required 30cN to be operated and led to one pellet each during training. For all other conditions, one of the levers was modified so as to double the amount of newtons required to operate it (60cN vs 30cN). This adjustment was done at the beginning of every session using a dynamometer (DM10, Nouvoutils). The hard lever’s side was randomized between rats, but was stable within rats. Animals had a maximum of 30mn to perform all 50 trials. Animals were considered ready for testing when rats performed > 70% of the free-choice trials within the final parameters. This last step of training consisted of three sessions of 50 trials each.

##### Habituation Actor & Victim

When actors had completed the training phase, the dyads were placed in the testing setup for 10min with house lights on, in order to habituate (i) the victim to the novel environment and (ii) the actor to the victim’s presence.

##### Exposure

When the actors were habituated to the presented of their partner, they were placed in the victim’s compartment (“Exposure”; Figure 1F) and were exposed to electric shocks [7]. This procedure consisted of a baseline (10min) followed by the delivery of four foot-shocks (0.8mA, 1s duration, 240-360s random inter-shock interval; yellow lightening in Figure 1). Actors were exposed alone in the setup. After the session, the actors were isolated in a small cage for 10 minutes to allow for the following animals to be placed in the exposure setup, so as to avoid the communication of stress to unexposed cage mates. For actors in the NonExposed condition (see below, and Table 2), the shockers were turned OFF during the Exposure session. For one group of rats, (Unfamiliar Victims condition), the exposure was performed in other cabinets using different odors and wall design [7], for historical reasons, as this condition was part of earlier experiments aimed to fine-tune our final experimental design.

##### Harm aversion testing

All dyads underwent four harm aversion testing sessions. The first session (“Baseline”; Figure 1E-F) took place on the day following the exposure, and was followed by 3 consecutive daily “Shock” sessions. Baseline and Shock sessions started with 4 forced trials (2 for each lever, pseudo-randomized) followed by 20 free choice (Figure 1E). This structure was constant for all conditions, except for the actors of the Unfamiliar Victims condition, where two baseline sessions followed by two shock sessions (each consisting of 4 forced and 8 free choice trials) were used for historical reasons. The forced trials were implemented to force the animals to sample each lever/outcome contingency at least twice per session. Failure to perform the forced trials restarted the trial (rats needed to perform all forced trials, i.e., were forced to sample each option twice). Failure to perform a free choice trial resulted in a failed trial, which incremented to the following trial. A timeout of 400s was used, after which the current trial was counted as failed (Figure 1G). Baseline and Shocks sessions were identical in all points except that during Shock sessions, pressing one of the two levers immediately delivered an electric foot-shock (0.8mA, 1s duration) to the victim (Figure 1F, “Shock”), while shocks were never delivered during baseline (Figure 1F, “Baseline”). Hence, the baseline sessions allowed to sample initial actor preferences for a given lever. While most actors preferred the easy lever in the baseline session (~75%), a subset preferred the hard lever. The lever that delivered a shock to the victim during harm aversion testing was determined based on preference levels at baseline (see also Supplementary Materials). This manipulation did not affect group differences in harm aversion behavior (Figure S1A). However additional training prior to harm aversion testing competed with harm aversion (Figure S1B). Animals were pre-fed 2h before beginning of each session with 66% of their daily food intake to avoid high levels of hunger which could have competed with harm aversion. The remaining 34% of the food allowance was provided after the experiment.

##### Experimental Conditions

Except for the familiarity control (see *“*Unfamiliar Victims*”* condition), actors and victims were housed in the same cage and were thus familiar with one another for 20 days before harm aversion testing started. In all cases animals were always tested with sex-matched conspecifics. Except for half of the dyads in the ContingentHarm condition (n = 12 actors), all tested dyads were males.

Different experimental conditions were tested in this study (Table 1 & 2). All conditions used different animals. Each actor was paired with a unique victim (sample size in Table 1 & 2 describes only the number of actors). The ContingentHarm condition (N = 12 male & N = 12 female actors) tested whether rats would decrease their preference for a lever upon its association with a conspecific’s distress. In this condition, the victim was immediately administered an electric shock every time the actor pressed the shock lever in the shock sessions, while rewards were administered 1s after lever press. Shock-lever association was deterministic, and the shock intensity was identical to the one used during the exposure session of the actor (1s duration, 0.8mA). In order to control for a spontaneous preference switch (i.e., not related to the shocks to conspecific), victims in the NoHarm condition (N = 14) were constantly isolated from shocks by using a plastic isolating floor during shock sessions. Hence, shocks were physically delivered to the grid floor, but never reached the victim. This was done to ensure that sounds associated with shock-delivery would be present in both conditions. In the RandomHarm condition (N = 8) the contingency between lever-presses and shocks was disturbed, while the number of shocks was conserved. This condition served as a control to ensure that higher social stress levels (caused by shocks delivered to the victim) cannot account for decreased preference for the shock lever. The 8 victims used in the RandomHarm condition received the same amount of shocks, and at the same trial number, as the 8 highest switcher animals in the ContingentHarm condition (determined using a permutation test; see “<u>*Statistical use of indexes*</u>”). Hence, stress levels in RandomHarm and ContingentHarm victims were comparable. Crucially, the shocks were <u>not</u> delivered at lever press, or contingent on which lever the actor presses, but at random intervals ranging between 3s and 8s during the inter-trial interval, which started after the actor exited the reward receptacle.

To explore the effects of familiarity, here we also present data from a pilot experiment in which the actors in the Unfamiliar Victims **c**ondition (N = 12) were paired with victims housed in a different cage, thus ensuring that the dyads were unfamiliar.

Additionally, we explored the effect of different cost modalities, i.e. differences between the two levers, on switching. To do so, we tested a 1vs2 pellets condition (1vs2Pellets; N = 7), a 1vs3 pellets condition (1vs3Pellets; N = 11) and we piloted a 0vs2 seconds condition (0vs2s; N = 5). In magnitude conditions, the levers required the same amount of force to be operated, but the shock lever delivered 2 (1vs2Pellets) and 3 (1vs3Pellets) pellets, while the no-shock lever delivered 1 pellet. In the delay condition, both levers required the same force to be operated and produced n = 1 pellet once pressed, but the shock lever delivered the pellet immediately (0s) while the no-shock lever delivered with pellet with a 2s delay. The difference in magnitude and delay was introduced on the baseline session, while training sessions featured same magnitude (1 pellet) and delays (1second) for each lever.

In order to test the effect of experience of shocks on the actor, actors in the NonExposed condition (N = 14) underwent the same treatment as the ContingentHarm animals except that they received no shock during the exposure session.

Finally, we investigated the effect of deactivating the ACC on harm aversion by using two additional groups of animals (Muscimol, N = 11; Saline, N =12). These animals underwent identical procedures as the ContingentHarm Condition, but were bilaterally implanted with internal cannulas targeting the ACC to inject muscimol or saline, respectively (see below).

##### Reversal control

A lack of behavioral flexibility could account for actors not switching away from the shock lever. To examine this possibility, all actors (except for 1vs2, 1vs3, 0vs2s and Unfamiliar Victim groups) underwent 3 consecutive reversal sessions (Figure 1E,F; Table 1 and 2) upon completion of the shock sessions. In reversal sessions, the lever which was not preferred (pressed <50% of the trials) during the last Shock session was associated with a higher reward (3 pellets) while the other lever still delivered the same amount (1 pellet). The actors went through three sessions of 24 trials (identical to the baseline session structure; Figure 1E) with the victim present in the adjacent compartment but without shocks involved during the session (Figure 1F, “Reversal”). Hence, the context was similar to the baseline and shock sessions, but the animal could reverse their previously acquired preferences to obtain more food.

##### Trial structure

Trial structure was identical across Baseline, Shock and Reversal sessions (Figure 1G). The trial started with illumination of the nose poke hole, inviting a nose poke. Actors were expected to perform a 400ms nose poke which triggered the presentation of the levers (only one lever in forced trials, both levers on free trials). Pressing either lever led to the delivery of sucrose pellets after 1s in adjacent reward receptacles (n=1 pellet for each lever in baseline and shock sessions for all conditions except 1v2 and 1v3, in which one lever lead to 1 pellet, and the other to 2 or 3 pellets, respectively; n= 1 and n = 3 pellets during reversal sessions), with the hopper illuminating for 0.5s upon delivery of the reward. During shock sessions (except for the NoHarm and RandomHarm condition) pressing the shock-lever additionally lead to an immediate 1s shock to the victim. An inter-trial interval of 10s started once the animal had exited the reward hole to consume the pellets, after which the nose poke was illuminated for the next trial to start. During baseline and shock sessions, a timeout of 400s was used to stop the current trial in case the animal did not perform a nose poke or did not press levers after the nose poke (Figure 1G; “Time Out”). This was deemed long enough to show a behavioral display of harm aversion (longer latency to consume food), while quantifying failed trials. In each trial, we collected two timing measurements happening after lever press: reward poke latency (from successful lever press to first entry in reward receptacle) and reward poke duration (time spent consuming the reward). Forced trials were systematically excluded from all analysis since (i) they did not reflect actual decisions from the actors and (ii) latencies might have been affected by acclimation time to the session.

#### Analysis and computations

In order to compare the effect of different manipulations at the group level, we use parametric frequentist (SPSS) and Bayesian statistics (JASP; https://jasp-stats.org/). We do so, because over the groups, lever preferences are approximately normally distributed, as assessed using the Shapiro Wilk test.

In order to explore how many individuals showed significant switching during the shock or reversal sessions, despite differences in baseline lever preferences, we also computed two indexes: the switching index, capturing changes in preference from baseline to shock sessions, and the reversal index, capturing changes in preference from shock to reversal sessions.

##### Switching Index

In order to quantify individual levels of switching despite differences in baseline preference, we computed a Switching Index (SI) for each rat, using the following equation, where S_baseline_ is the proportion of shock lever presses during baseline, and S_shock_ the average proportion of shock lever presses over all three shock sessions.

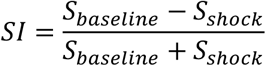

The SI*s* are individual values that range between [-1/3;1] and quantify the strength of switching between baseline and shock sessions. The distribution of potential SIs can be found in Figure 1F. Possible values for S_baseline_ were sampled uniformly from the interval [0.5;1], given that S_baseline_ cannot be below 0.5, because the shock lever is the lever preferred at baseline, by definition. Possible values for S_shock_ were sampled from the interval [0;1].

##### Reversal Index

In order to quantify the strength of reversal learning, we computed a Reversal Index (RI) for each individual. In reversal sessions, the least preferred lever during shock sessions was now associated with n = 3 pellets, whereas the remaining lever produced n = 1 pellets. The RI was computed as follow:

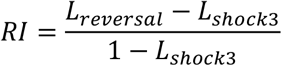

Where L_reversal_ is the proportion of choice for the lever producing 3 pellets averaged over all reversal sessions, and L_shock3_ that during the last shock session. This normalizes the change in preference by the maximum potential change possible in preference. The RIs are individual values that range between −1 and 1 and quantify the preference for the 3 pellets option. The distribution of potential RIs can be found in Figure 1F. Possible values L_shock3_ were sampled uniformly from [0;0.5], because the 3 pellet lever is by definition that non-preferred in shock3. Possible values for L_reversal_ from [0;1].

##### Statistical use of indexes

The equations used for SI and RI create skewed distributions (Figure 1F). To overcome these issues, we used non-parametric permutation analysis to detect animals that showed significant switching. We generated, for each animal separately, a distribution of permuted indexes, computed by shuffling the choices (with replacement, N = 10.000 times) across baseline and shock sessions (SI), or between shock3 sessions and reversal sessions (RI). We then compared actual indexes to the 95 % confidence interval (CI) of this randomized distribution of social bias scores. We provide the individual CI’s lower and upper bound in the supplementary materials; Table S1-3. In addition, to explore whether the observed effects could be due to the bias introduced by selecting the shock and reversal levers as those above and below 50%, respectively, we also concentrate analyses on comparing choices across experimental and control conditions, which suffer from the same bias.

#### Additional Behavioral Analysis

##### Video rating

To get a more detailed view of the rats’ behavior, the videos recorded were manually scored with the use of the program Solomon Coder beta 17.03.22. (https://solomoncoder.com/) The behaviors scored for the actor consisted of: rearing, grooming, freezing, and jumping.

##### Data analysis

MedAssociates and Solomon behavioral logfiles data extraction was performed with the use of MATLAB R2017b. Further statistical analyses were performed with IBM SPSS Statistics version 25.00 for Windows. Data are expressed as means ∓ standard errors of the means (sem). The significance level was set at *p* < .05. Unless specified otherwise, all the displayed results correspond to behavioral data of the actor. The choice preference for the shock lever over sessions was analyzed with a repeated measures analysis of variance (*rm*ANOVA) with session as a within-subjects factor (baseline and shock 1-3). The above stated analysis was also used for reversal data and freezing behaviour. Post-hoc pairwise comparisons are reported in the figures. Pairwise comparisons consisted of paired sample t-tests when comparing different epochs of the same group, and independent sample t-tests when comparing different groups within an epoch. Given the low sample size of the Saline and Muscimol groups in the reversal sessions, we used the non-parametric equivalent (Wilcoxon tests). Comparisons were adjusted for multiple comparisons using the Benjamini & Hochberg correction (i.e., false discovery rate) using the R statistical package (R stats package). Latencies for lever press, reward poke, nose poke and reward poke duration were log-transformed to meet parametric testing assumptions.

##### Interpretation of Bayes factor

We computed Bayesian statistics (JASP) in order to quantify the evidence for absence of effect when frequentist statistics showed non-significant effects (*p* > .05). Conventionally, the generated quantity (Bayes Factor in favor of H1; BF_10_) should be interpreted as a relative plausibility of the H_1_ hypothesis over the H_0_ hypothesis. If BF_10_ > 3, there is moderate evidence in favor of H_1_. If BF_10_ < 1/3, there is moderate evidence for H_0_ (evidence for the absence of effect). If 1/3 < BF_10_ < 1, no strong conclusion should be drawn based on this data regarding the existence or absence of effect, but the relative plausibility of both hypotheses can be appreciated. When using two-tailed hypothesis, the BF is written as BF10, when using one-tailed hypothesis of the experimental condition having lever-preferences lower than the control condition, as BF-0, as recommended in JASP.

#### Surgery and cannulation

After training was completed, rats in the muscimol and saline conditions (N_total_ = 27) underwent a surgical procedure for the bilateral implantation of cannulas targeting the ACC. Body temperature and other physiological parameters were monitored throughout the surgery. Rats were anaesthetized using isoflurane (5% induction, 2/2.5% maintenance), and prepared for surgery by weighing, and shaving the head. Once animals were placed on a stereotaxic apparatus, the incision area was cleaned with alcohol/betadine and sprayed with 10% xylocaine (lidocaine, spray) used as a local anesthetic. Two holes were drilled to bilaterally implant with stainless steel cannulas targeting the ACC (Plastic One, C313G/Spc 3.5mm). The cannulas were placed at the following coordinates: AP, + 1.17 mm; ML, ∓1.16 mm; DV, +1.8 mm (AP & ML from bregma; DV from the surface of the skull; [15]). All coordinates were taken based on [32]. Three additional holes were drilled around the implanted cannula to place steel screws, which were later used to anchor the cannula to the skull using dental cement (Prestige Dental, Super Bond C&B Kit, UK). To minimize the damage to the cannula, they were covered with a protection cap (Plastic One, C313DC/1/Spc 3.5mm). After the surgery, an analgesic/anti-inflammatory drug was delivered for pain relief (Metacam, 2 mg/kg, sc) and 0.5ml of saline sc was given for rehydration. Animals were then placed in an incubator until they woke up. The animals were housed individually for 10 days to recover from the surgery. To monitor any possibility of discomfort or pain and to ensure that the animals were having a proper recovery process, the appearance, behavior, state of the incision (wound healing), recovery process and weight were monitored daily for 10 days after the surgery. Two animals died during surgery. Post-mortem dissection revealed a small heart lumen which might have rendered animals more sensitive to isofluorane anesthesia. One additional animal showed abnormal weight loss and aberrant behavior, and was euthanized two days after surgery (euthanized in CO_2_ chambers with initial 40% O_2_ mixed with 60% CO_2_ until animals were in deep sleep as verified through paw reflexes, then switched to 100% CO2 for at least 15 minutes). For the remaining 24 animals following the surgery, all rats lost minimal or had no weight loss, and the animals that lost some weight returned to normal weight gain following 2 to 3 days after the surgery. In addition, all these animals showed normal behavior, prompt recovery and healthy wound healing following the surgery. After 10 days of recovery, animals were tested two days in the operant box (same design as training, step 3) to ensure that the implanted cannula did not affect food access, and that operant behavior was unaffected by the intervention.

#### Humane endpoints

The humane endpoints were as follows:

1. *Insufficient recovery after surgery:* It was considered if animal showed permanent weight loss. The threshold was set to a 15% weight loss after surgery monitored during 10 days.

2. *Infection:* Although we always perform the surgeries in sterile conditions, there was a small possibility of infection around the wound area. Visible signs of pathogenesis were monitored. The following were considered as signs of unhealthy state of the animal: aberrant behaviour, shock, dehydration, weight loss, nose and mouth discharge, bleeding, fits/seizures, diarrhea.

#### Infusion of saline and muscimol

Actor rats in the Muscimol condition were given micro-injections of the GABAa receptor agonist muscimol before the start of the behavioural testing in baseline and shock sessions. Muscimol (Sigma Aldrich, M1523) was dissolved in sterile phosphate buffered saline to obtain a final concentration of 0.1ug/ul. Infusions (0.5ul per hemisphere) were made bilaterally under isoflurane anesthesia, which lasted on average 7min. This time included 2min of induction, 2.5min of infusion time (0.25 ul/mn infusion rate) and 2.5min of diffusion time (with infusion needle left in place). Infusions were made in both hemispheres simultaneously using an infusion pump (Syringe Pump PHD Ultra Infuse, Harvard Apparatus) equipped with a micro-dialysis rack for 4 syringes (Harvard Apparatus). In the Saline condition, the exact same procedure was followed except that phosphate buffered saline (0.9%) was used without muscimol. From the end of infusion, a delay of 20min was implemented before the start of the sessions to allow full effect of muscimol on brain tissue.

#### Histology

Upon completion of the reversal sessions, animals were anaesthetized using isoflurane (5% induction). Depth of anesthesia was checked by verifying the paw reflexes, after which animals were perfused transcardially using 0.01M phosphate buffer (PBS, 0.1M, pH = 7.4) for three minutes followed by a fixating solution of paraformaldehyde (PFA, 4%) for 5min. Brains were immediately removed and stored in PFA solution for ten days at a temperature of 5°C. Coronal sections (50 um) of the ACC were obtained using a vibratome (Leica VT1000S, Germany) and mounted for histological examination. Finally, injection sites were mapped using a microscope (Zeiss Axioplan 2, Germany) and the a rat atlas [33] with standardized coordinates. Due to methodological issues during brain extraction and slicing, this analysis resulted in N = 9 brains mounted in each group (muscimol and saline). Coordinates shown in Figure 5 were the result of combining the evidence about location from both hemispheres. Victims were euthanized in CO_2_ chambers with initial 40% O_2_ mixed with 60% CO_2_ until animals were in deep sleep, as verified through paw reflexes, then switched to 100% CO2 for at least 15min. One animal was discarded due to clogged internal cannula during infusion in the baseline session. The remaining animals (muscimol, N = 11; saline, N = 12) were used for behavioral analysis.

#### Data Availability

Choice data is provided in data.xlsx. All other data will be made available upon publication.

## Author Contributions

J.H-L. Acquired funding, conceived the study, conducted all experiments, analyzed the data and wrote the manuscript.

A.T.A. Acquired data for the ACC deactivation and the effect of prior exposure, analyzed the data for these sections and provided comments on the manuscript.

V.G. Acquired funding, conceived the study, analyzed some of the data and wrote the manuscript.

C.K. Acquired funding, conceived the study, analyzed some of the data and wrote the manuscript.

## Acknowledgments

J.H-L is supported by an Individual Marie Curie Fellowship (SocioRats, #745885). C.K is supported by a VICI grant attributed to C.K. V.G. is supported by a VIDI grant (#XXX) attributed to V.G. C.K., V.G. and J.H-L. We thank Efe Soyman and Steven Voges for valuable comments on the manuscript. We thank Susan van der Boogard for her help with the data acquisition, as well as Laura Stolk for her assistance in histological verifications in cannulas implantation. We thank the Mechatronics department of the Netherlands Institute for Neuroscience for their help in building the experimental setups. We thank animal care takers of the Netherlands Institute for Neuroscience for valuable support with animal care.

---

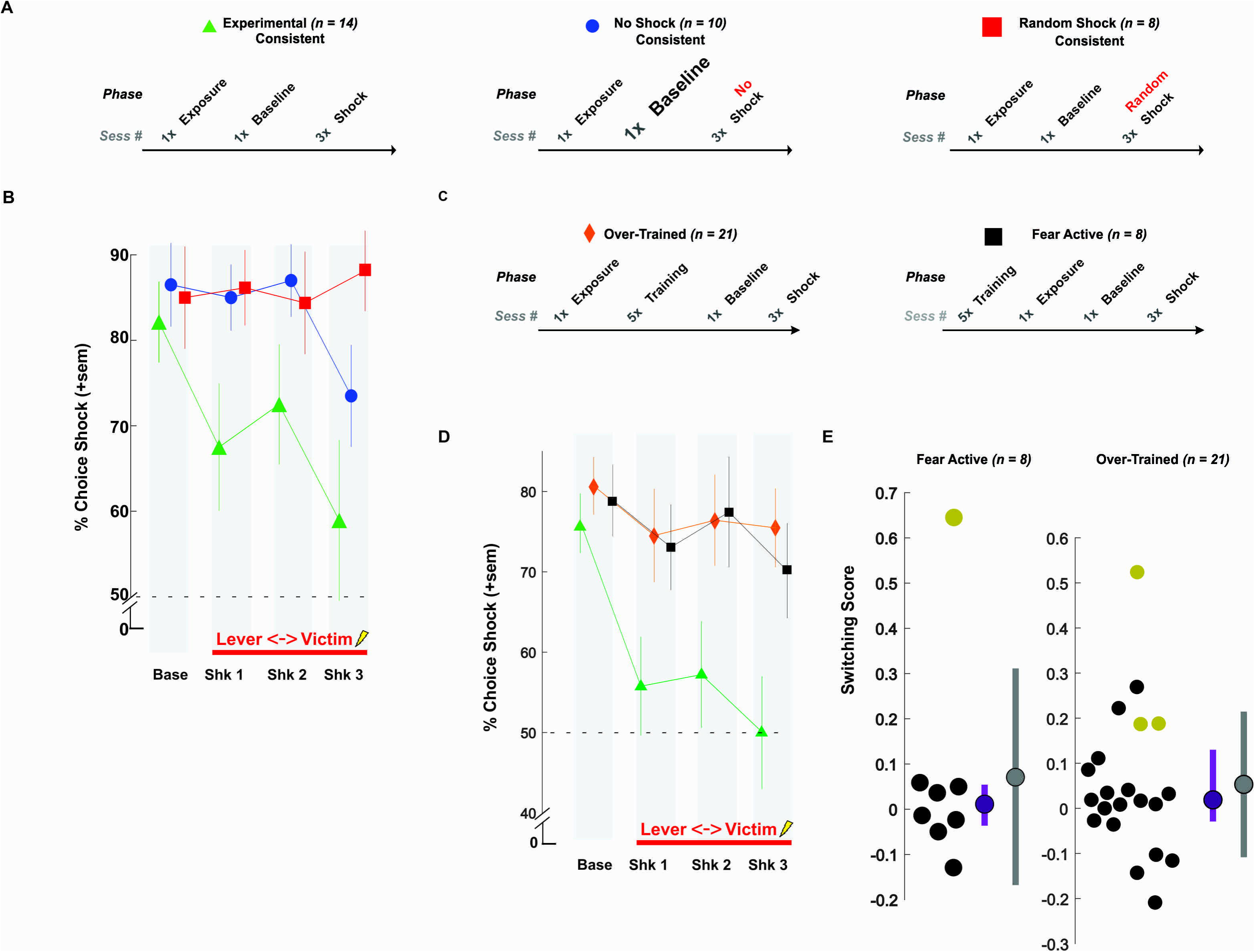

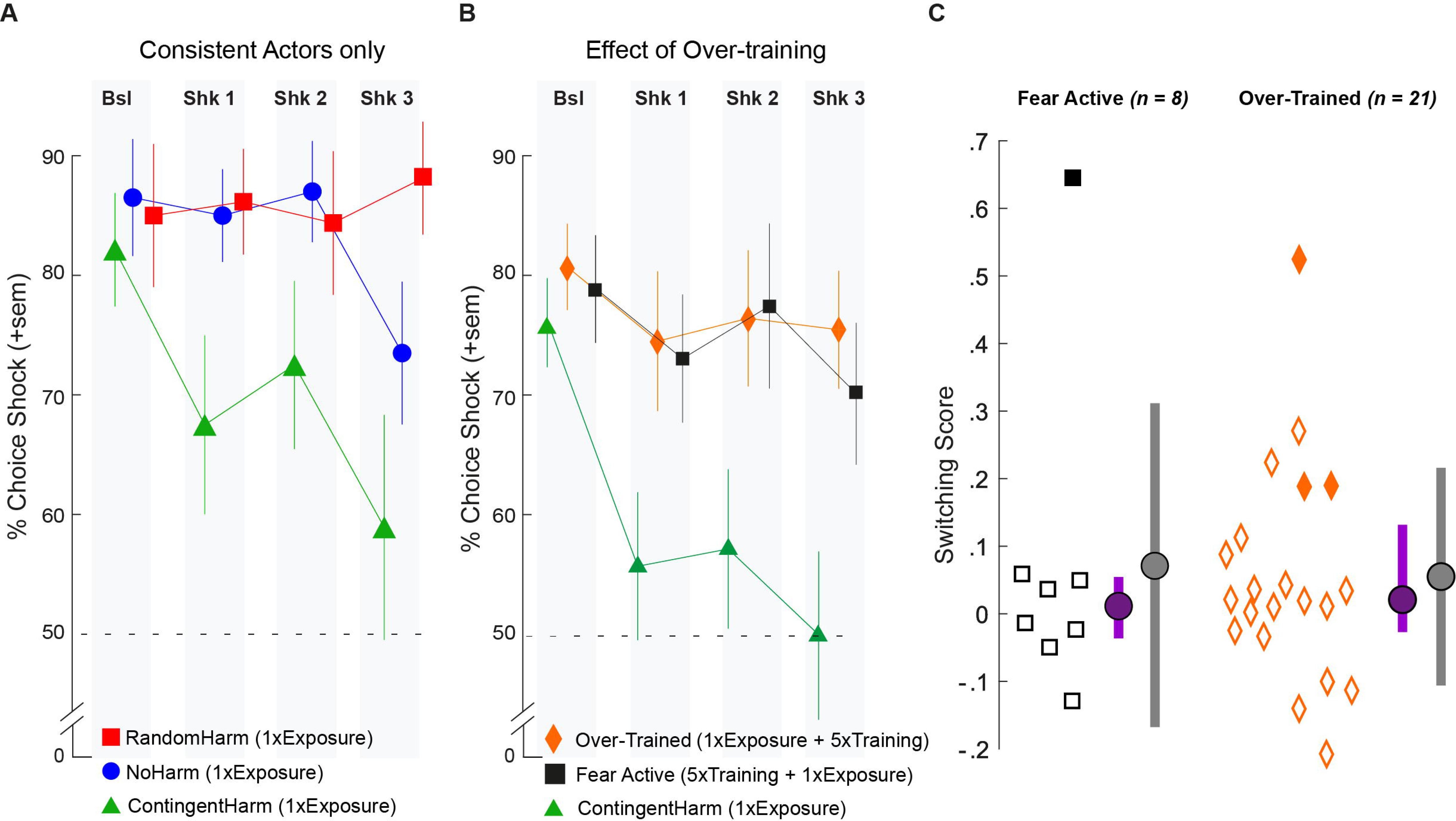

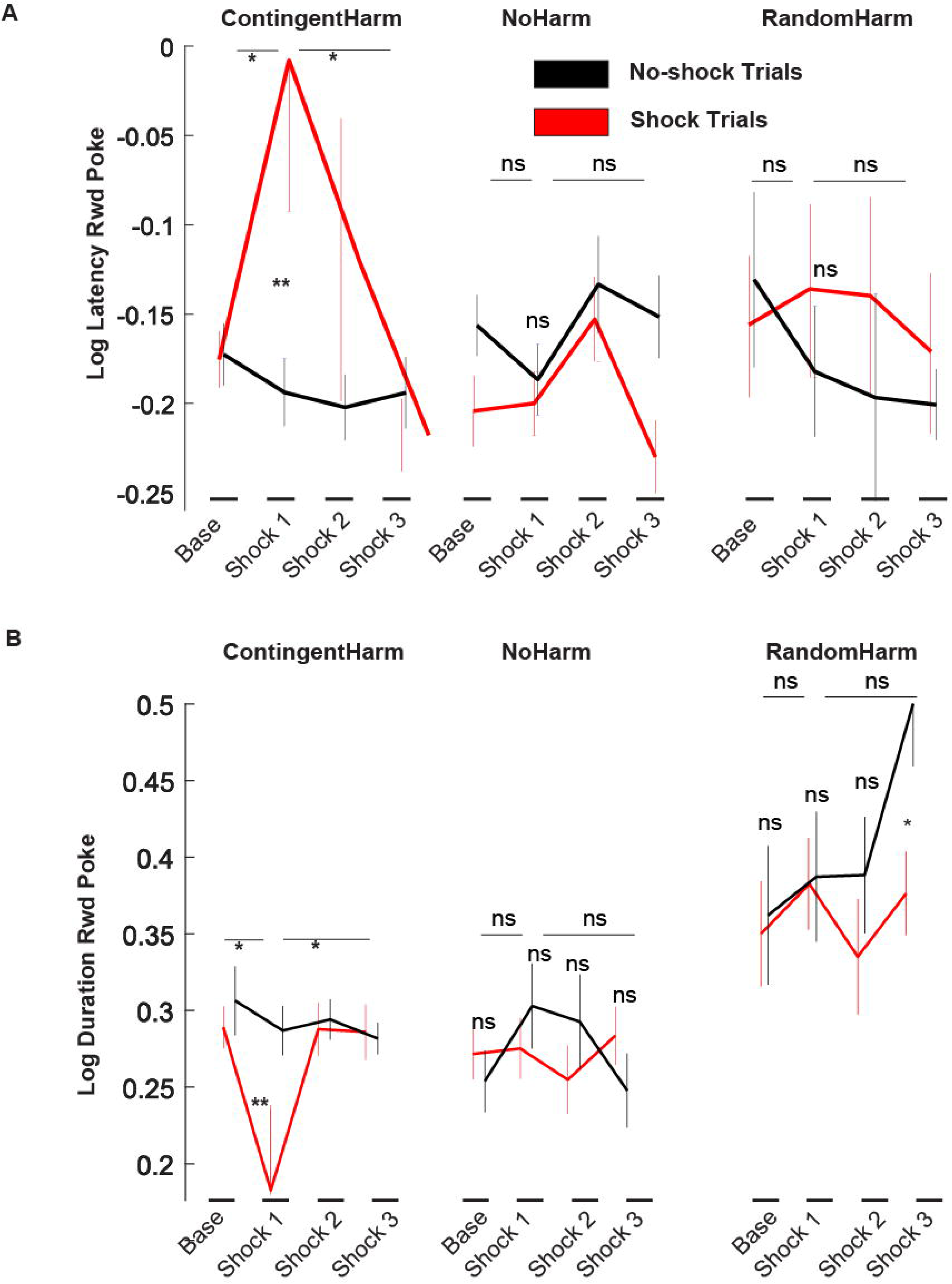

