## Supplemental Materials for "Harm to others acts as a cingulate dependent negative reinforcer in rat"

**Lead Author: Christian Keysers**

**Supplementary Methods**

*Determining shock lever:* The lever that delivered a shock to the victim during prosocial testing was determined based on preference levels at baseline. Hence, the lever that was chosen > 50% of the trials during baseline session was associated with a shock during the subsequent three shock sessions. This manipulation might have induced a bias, since it both artificially increased (i) baseline levels and (ii) the likelihood of obtaining a decrease of preference from baseline. However, since all groups were subjected to the same manipulation, group differences are statistically valid. An alternative way to determine which lever to use as a shock-lever would be to select the lever that is preferred at the end of the training session before the baseline. To test if such a selection would alter results, we repeated the analysis only for those animals that had the same preference during training and baseline and for whom we would thus have the same results if we had selected the shock lever based on training instead of baseline trials. We call these animals ‘consistent’. The analysis ran with the remaining, consistent animals (Figure S1A) again showed a significant effect of session (*F_(3,87)_ = 3.50, p = .019, η^2^ = 0.11*) and interaction (*F_(6,87)_ = 2.47, p = .03, η^2^ = 0.15*), suggesting that including inconsistent actors does not artificially drive the prosocial effect.

In early experiments, we had aimed to avoid any switch in preference between training and baseline by including more training trials until the preference was stable. In these early groups of animals (“Over-Trained Condition”, n = 21) actors received additional training sessions (5 sessions on average; Figure S2B), and remained in training until the proportion of choice for a given lever was higher than 80% during two consecutive training sessions (instead of one session > 70% correct trials with a nosepoke duration of 400ms, regardless of preference levels, in the groups in the main paper). In the Over-trained condition, we also performed the exposure before the training. In that group we failed to find significant switching when associating the preferred lever during the last training session with shocks in the shock sessions (Figure S1D, rmANOVA, 4 sessions, *F_(3,60)_ = .97, p =.41, BF_10_ =.18*). In order to isolate whether the lack of switching was due to overtraining or to performing the exposure further away in time from the shock trials, an additional group of animals (“Fear Active”, n = 8) was trained like the Over-Trained animals, but the exposure was performed after the additional training sessions as in our main experiments(Figure S1B). This group also did not show significant switching (4 Session rmANOVA, *F_(3,18)_ =.9, p = .46, BF_10_ = .36*). To verify that overtraining reduces switching, we performed a 3 condition (ContingentHarm vs Over-Trained vs Fear Active) x 4 sessions *rm*ANOVA, which revealed a significant effect of session (*F_(3,144)_ = 6.96, p < .001, η^2^ = 0.13*) and significant interaction session*condition (*F_(6,144)_ = 2.65, p = .027, η^2^ = 0.10*), which was driven by the animals in the ContingentHarm group, while no significant session differences were found in the OT and FA groups (all *p* > .05). Single subject analysis on the switching index revealed that only 1/8 (12%) and 3/31(14%) in the Fear Active and Over-Trained group, respectively, showed significant switching, three times fewer than the 38% switchers in the ContingentHarm group (Figure S1C). These results suggest that highly robust baseline preferences obtained with extensive training work against harm aversion in rats.

*Individual analysis:* A permutation test was used to determine significant switchers in each group. The 95% chance interval of the SI obtained by permutation are depicted in the tables below.

| **Rat #** | 1 | 2 | 3 | 4 | 5 | 6 | 7 | 8 | 9 | 10 | 11 | 12 |
| --- | --- | --- | --- | --- | --- | --- | --- | --- | --- | --- | --- | --- |
| **Low bound** | -0,52 | -0,31 | -0,33 | -0,48 | -0,15 | -0,18 | -0,26 | -0,47 | -0,11 | -0,13 | -0,03 | -0,37 |
| **High bound** | 0,38 | 0,24 | 0,22 | 0,37 | 0,13 | 0,17 | 0,18 | 0,32 | 0,08 | 0,12 | 0,08 | 0,27 |
| **Rat #** | 13 | 14 | 15 | 16 | 17 | 18 | 19 | 20 | 21 | 22 | 23 | 24 |
| **Low bound** | -0,12 | -0,68 | -0,19 | -0,04 | -0,20 | -0,03 | -0,45 | -0,12 | -0,45 | -0,28 | -0,21 | -0,21 |
| **High bound** | 0,09 | 0,36 | 0,19 | 0,03 | 0,15 | 0,08 | 0,28 | 0,09 | 0,41 | 0,22 | 0,13 | 0,17 |

***Table S1 | Chance interval bounds generated by permutation test for ContingentHarm group (n = 24)***

| **Rat #** | 1 | 2 | 3 | 4 | 5 | 6 | 7 |
| --- | --- | --- | --- | --- | --- | --- | --- |
| **Low bound** | -0,18 | -0,15 | -0,03 | -0,18 | -0,16 | -0,24 | -0,11 |
| **High bound** | 0,17 | 0,11 | 0,01 | 0,15 | 0,14 | 0,17 | 0,10 |
| **Rat #** | 8 | 9 | 10 | 11 | 12 | 13 | 14 |
| **Low bound** | -0,09 | -0,06 | -0,25 | -0,15 | -0,29 | -0,14 | -0,07 |
| **High bound** | 0,10 | 0,04 | 0,20 | 0,11 | 0,20 | 0,11 | 0,03 |

***Table S2 | Chance interval bounds generated by permutation test for NoHarm group (n = 14)***

| **Rat #** | 1 | 2 | 3 | 4 | 5 | 6 | 7 | 8 |
| --- | --- | --- | --- | --- | --- | --- | --- | --- |
| **Low bound** | -0,11 | -0,03 | -0,14 | -0,14 | -0,12 | -0,07 | -0,21 | 0 |
| **High bound** | 0,09 | 0,01 | 0,13 | 0,11 | 0,08 | 0,03 | 0,13 | 0 |

***Table S3 | Chance interval bounds generated by permutation test for RandomHarm group (n = 8)***. Animals #8 had constant pressing of one lever, which provided no power to compute a CI for this animal.

**Supplementary Figures (numbered in relation to main figures)**

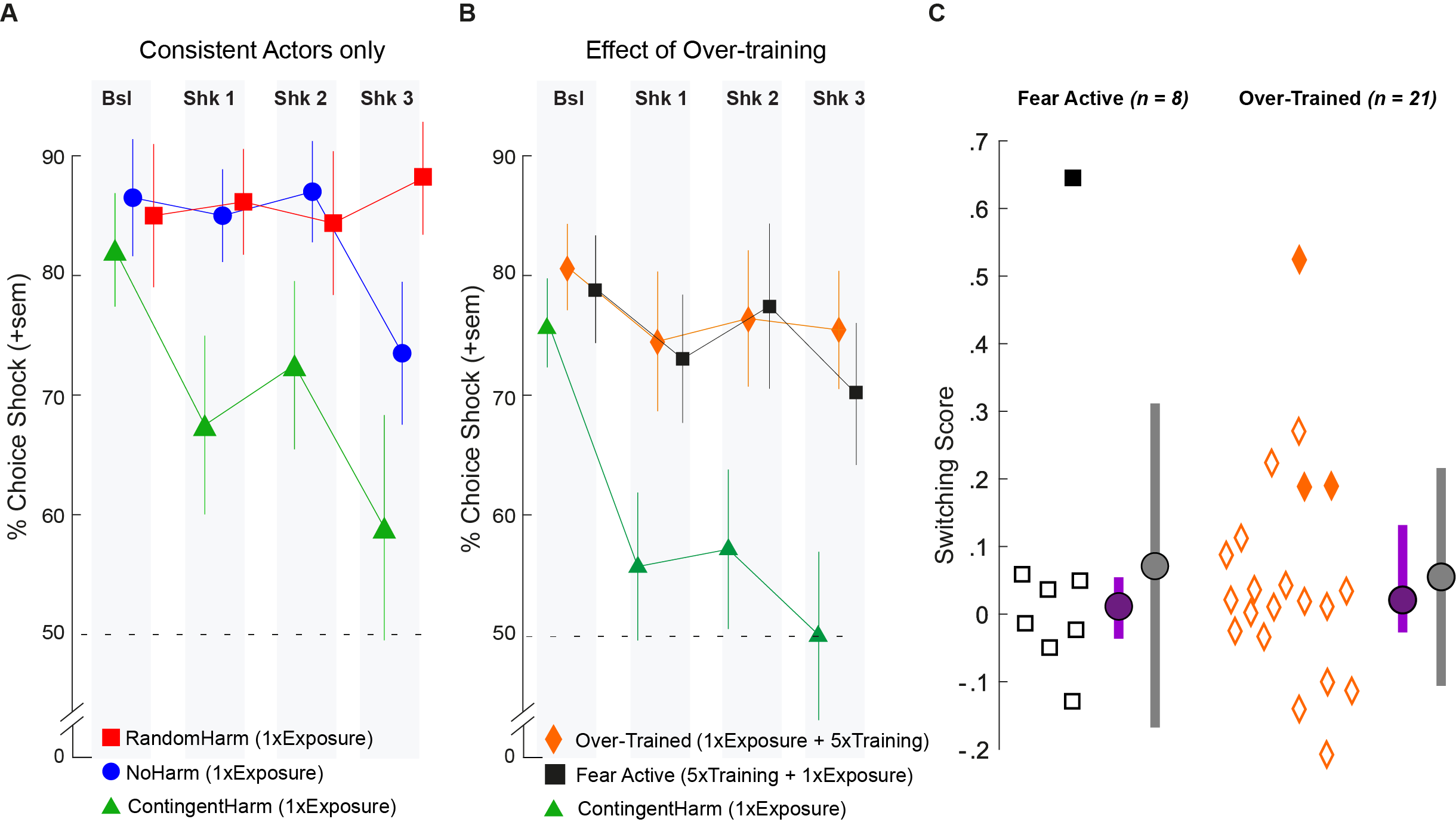

***Figure S1 | Effects of Consistency and Overtraining (A)*** *Percent choice for the shock lever for consistent actors only (i.e., actors that had higher proportion of choice for the same lever across training and baseline sessions).* ***(B)*** *Percent choice for the shock lever in the Over-Trained and Fear Active condition that had received 5 additional training sessions either after or before exposure. In both panels. Data is shown as mean +/- s.e.m (C) Switching scores for Fear Active and Over-Trained conditions. Empty symbols indicate non-switchers, filled symbols switchers. Purple dot and line are the distribution’s median and the 25 and 75 % percentile values, respectively. Grey dot and line are the distribution’s mean and standard deviation, respectively. Related to Figure 2*

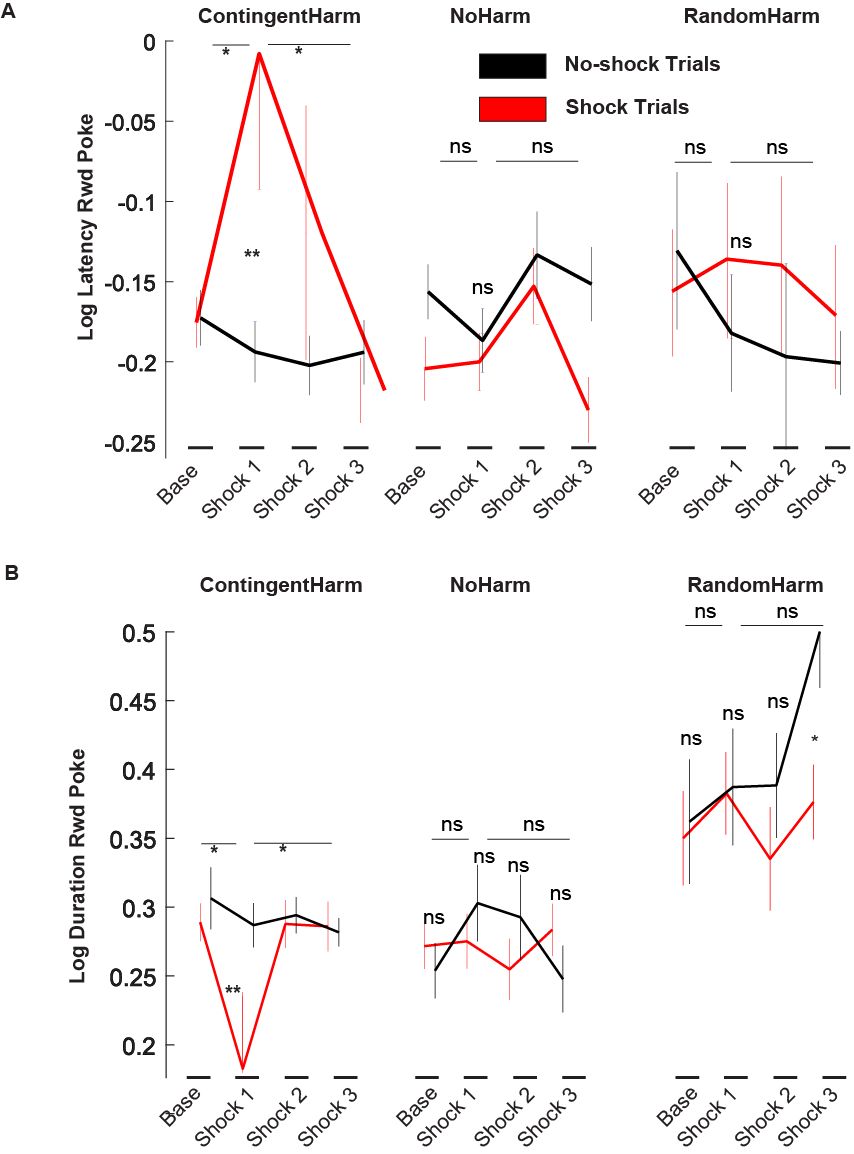

***Figure S2 | Changes in consummatory behavior following shocks. (A)*** *Reward poke latency for shock (red) and no-shock trials (black) in the ContingentHarm, NoHarm and RandomHarm group. In the ContingentHarm group, we found a significant effect of session (F(3,39) = 2.97; p = .043; η2 = 0.19) and a significant interaction (session*trial[Shock or NoShock]; F(3,39) = 3.22; p = .033; η2 = 0.20). Post-hoc pairwise comparisons revealed that the log-latency to reward poke after pressing the shock lever increased dramatically in the first shock session compared to baseline (two-tailed; fdr corrected; baseline vs Shock 1: t(22) = 2.27; p = .03; CI = [-.01 ; .34]), and returned back to baseline levels in the following two shock sessions (Shock 1 vs Shock 2: t(22) = 1.95; p = .19; CI = [.02 ; .36]; Shock 1 vs Shock 3: t(20) = 2.28; p = .01; CI = [.02 ; .36]). Importantly, the log-latency increased only when animals pressed the shock lever (two-tailed; Shock 1; trial[shock vs no-shock]: t(20) = 2.04; p = .006; CI = [-.32 ; -.06] ; all other comparisons: p > .05). No significant effects on log-latency to reward poke were found in the NoHarm and RandomHarm groups (all p > .05).* ***(B)*** *Similar effects were found on reward poke durations, i.e., the time spent consuming the reward (session: F(3,39) = 2.49, p = .07, η2 = 0.16; session*lever: F(3,39) = 3.06, p = .04, η2 = 0.19). Animals in the ContingentHarm group showed a significant decrease in reward poke log-duration from baseline to first shock session (two-tailed; fdr corrected; baseline vs Shock 1: t(21) = 2.08, p = .01, CI = [.01 ; 0.21]), which increased in the second and third shock sessions (Shock 1 vs Shock 2: t(20) = -2.07, p = .01, CI = [-.13 ; .01]; Shock 1 vs Shock 3: t(19) = -2.57, p = .01, CI = [-.14 ; -.01]). Importantly, the log-duration decreased only when animals pressed the shock lever (two-tailed; Shock 1; trial[shock vs no-shock]: t(20) = -1.76, p = .005, CI = [.03 ; .18] ; all other comparisons: p > .05). No significant effects were found in the NoHarm group (all p > .05; Figure S3). Animals in the RandomHarm groups showed a decrease in reward poke log-duration across sessions in the shock trials (Figure S3), which reflected the victim’s shock timing (i.e., delivered during the ITI in the RandomHarm group), which interrupted reward poke log-duration in shock, but not in no-shock trials. Related to Figure 3*
